# Divergent scalp-to-region distance alteration patterns in autism spectrum disorders, Parkinson’s disease and Alzheimer’s disease

**DOI:** 10.64898/2026.05.14.725296

**Authors:** Liqin Yang, Junhao Zhang, Junlong Wang, Hsin-Hsiung Huang, Hongbin Han, Daniel Razansky, Alzheimer’s Disease Neuroimaging Initiative, Axel Rominger, Jie Lu, Ruiqing Ni

## Abstract

Brain stimulation is increasingly recognized as an effective and important therapeutic intervention for many brain diseases. Distance between the scalp and other brain regions is a pivotal variable for neurostimulation planning and the development of new techniques, but alterations in the distance between the scalp and other regions in brain diseases are largely unknown. In this study, we developed an automatic pipeline to calculate scalp-to-region distance (SRD) values from T1 MR images and applied it to a total of 1382 participants, including patients with autism spectrum disorder (ASD), Parkinson’s disease (PD), Alzheimer’s disease (AD), and cognitively normal controls (CNs). Cloud points were uniformly sampled on the automatically extracted scalp surface and cortex surface, on which the point-wise distance maps were generated. The brain was then coregistered with the BCI-DNI atlas, and SRD value for each brain region was extracted. Analysis of covariance (ANCOVA) was performed for SRD in each brain region, with age and sex as covariates. Compared with CNs, ASD patients showed widespread SRD decreases across the brain with prominent involvement of the frontal lobe, especially the orbitofrontal cortex and adjacent regions. In contrast, in AD patients, significantly increased SRD values were observed in various regions of the frontal gyrus. No significant SRD alteration was found in PD patients after correction. The automatic SRD calculation pipeline and the different patterns of SRD alterations in these diseases might be helpful for future neurostimulation planning in clinical practice.

**Highlights:**

1. Automatic pipeline enables scalp-to-region distance (SRD) measurement, facilitates brain stimulation planning.
2. ASD patients show widespread SRD decreases, especially in the orbitofrontal cortex and adjacent regions.
3. AD patients present increased SRD in the frontal gyrus and decreased SRD in the parahippocampal gyrus.

## 1. Introduction

Brain stimulation is increasingly recognized as an effective and important therapeutic intervention for many brain diseases. In addition to invasive techniques such as deep brain stimulation (DBS) [1], non-invasive neurostimulation techniques, including transcranial electrical stimulation (tES) techniques such as transcranial direct current stimulation (tDCS) [2], temporal interference stimulation (TI) [3, 4], transcranial magnetic stimulation (TMS) [5, 6], transcranial ultrasound stimulation (TUS) [7, 8], and transcranial pulse stimulation (TPS) [8, 9] etc., are currently under rapid development and towards clincal application. Neurostimulation planning is important for precise individualized treatment in clinical practice, involving the determination of both the optimal stimulation target and corresponding parameters such as orientation, trajectory and dosage[10]. Distance between the scalp and targeted brain regions is an essential variable for planning neurostimulation and for optical imaging of the brain [11-13]. Notably, noninvasive neurostimulation techniques rely on the transversing of signals between the resources placed on the scalp and the targeted brain regions[14, 15]. Signals based on electromagnetic fields, such as those in transcranial magnetic stimulation, decay with distance[16]. The amount of intensity and the energy needed for TMS to induce motor movement (called the motor threshold) has been demonstrated to depend on the distance from the coil or scalp to the underlying cortex[17, 18]. In addition to inter-individual variability, disease-level characteristics of both the scalp and the brain represent critical factors for the development of novel neurostimulation targets and treatment planning. Inspired by earlier group-level discoveries of disease-related brain alterations, disease-specific targets have been identified and implemented for various disorders, such as the involvement of the subgenual anterior cingulate cortex in depressive disorder[19]. In contrast, potential scalp-related alterations in diseased populations have received relatively little attention. In light of the above, it is important to comprehensively investigate the scalp-to-(brain)region distance of patients before brain stimulation.

Several scalp-to-cortex distance (SCD) analysis tools based on magnetic resonance imaging (MRI) data have been developed in recent years, but research on SCDs in different brain diseases is limited. SCD values can be calculated by manually selecting points or regions of interest and then measuring a line or sampling [17, 20, 21]. In the studies of Lu et al. [22-24], they selected brain region points according to the Montreal Neurological Institute (MNI) coordinates and their corresponding locations on the scalp by cursor in a neuronavigation system and then adjusted the TMS coil. Using this method, they analyzed the dorsolateral profrontal cortex (DLPFC), a typical target region for TMS treatment, in patients with dementia[22], Parkinson’s disease [23] and mild cognitive impairment converters[24], with a limited sample size (<50 patients). While these manual or semi-automated methods are feasible for individual-level or small-sample analyses focusing on specific regions, their application to multiple regions and larger cohorts is hindered by inevitable challenges involving reproducibility and time efficiency. Van Hoornweder et al. [25] developed a SCD tissue analysis toolbox to analyze data from 250 younger and older adults and reported differences related to age, sex and channel regions. Recently, we proposed a fully automated open-source pipeline to efficiently generate SCD maps with densely sampled points covering the entire skull and cortex surface[26]. We applied this pipeline to 407 cognitively normal elderly individuals and revealed its association with age and sex. To date, the SCDs of various brain diseases have rarely been studied, especially at multi-region level and in large datasets.

In this study, we developed an automatic pipeline to calculate regional SCD values, i.e., scalp-to-region distance (SRD) values, and applied it to cohorts of 1382 participants, including cognitively normal controls (CNs), patients with autism spectrum disorder (ASD), Parkinson’s disease (PD) and Alzheimer’s disease (AD). We hypothesized that the three diseases should have disease-specific SRD alteration patterns that are helpful for planning their brain stimulation in the future.

## 2. Methods

### 2.1 MRI data

Data used in the preparation of this article were obtained on January 2023 from the Alzheimer’s Disease Neuroimaging Initiative (ADNI) (adni.loni.usc.edu) [27, 28], the Parkinson’s Progression Markers Initiative (PPMI) database (www.ppmi-info.org/access-data-specimens/download-data) (RRID:SCR_006431) [29] and the Autism Brain Imaging Data Exchange (ABIDE) database [30, 31]. The ADNI was launched in 2003 as a public□private partnership led by Principal Investigator Michael W. Weiner, MD. The primary goal of the ADNI has been to test whether serial MRI, positron emission tomography (PET), other biological markers, and clinical and neuropsychological assessments can be combined to measure the progression of mild cognitive impairment (MCI) and early AD. The PPMI is an observational study launched in 2010 to identify and validate biomarkers of Parkinson’s disease progression, including clinical, imaging, and biospecimen measures. ABIDE was established to aggregate and share MRI data across multiple international sites for the purpose of large-scale evaluation of the intrinsic brain architecture in individuals with ASD. The demographic information of the participants included in this study is presented in **Table 1**. T1W MR images of each participant obtained with 3T MRI scanners were used. For participants with longitudinal imaging data, only images at baseline were selected. These anatomical MR images, although collected from different cohorts and institutes, have similar image resolution (voxel size: ∼ (1–1.25) × (1–1.25) × (1–1.5) mm^3^). The detailed MRI imaging protocols of each cohort and institute can be found on the respective website.

**Table 1.** Demographic data.

| **ABIDE** | **CN** | **ASD** | **P value** |
| --- | --- | --- | --- |
| N | 381 | 313 |  |
| Sex (M/F) | 337/44 | 258/55 | 0.024 |
| Age (years) | 15.88 ± 6.34 | 15.7 ± 5.69 | 0.6883 |
| **PPMI** | **CN** | **PD** | **P value** |
| N | 84 | 225 |  |
| Sex (M/F) | 55/29 | 144/81 | 0.8095 |
| Age (years) | 59.5 ± 11.62 | 61.73 ± 9.49 | 0.085 |
| **ADNI** | **CN** | **AD** | **P value** |
| N | 224 | 155 |  |
| Sex (M/F) | 114/110 | 75/80 | 0.6315 |
| Age (years) | 76.04 ± 6.42 | 74.66 ± 7.43 | 0.0534 |

CN, cognitively normal; ASD, autism spectrum disorder; PD, Parkinson's disease; AD, Alzheimer's disease; Age data are presented as the mean ± standard deviation.

### 2.2 MRI data analysis pipeline

We updated the previously developed open-source pipeline BrainCalculator [26] by incorporating an atlas registration module to compute the SRD values for different regions (**Fig. 1**). Taking a T1-weighted brain MR image as input (**Fig. 1a**), the BrainSuite (BS) 21a [32] skullfinder toolbox was used to segment the scalp (**Fig. 1b**) and rough cortex surfaces based on a sequence of edge detection and mathematical morphology operators [33]. Next, the image was further processed by a series of topology correction and masking algorithms to extract the pial surface (the outer cortex surface) of the brain [34] (**Fig. 1c**). After the scalp surface and cortex surface were extracted, 5× 10^5^ points were uniformly sampled on the two surfaces (**Fig. 1d&e**). For each point on the cortex surface, the scalp-to-cortex distance was calculated as the shortest 3D Euclidean distance between all the scalp points and that cortex point (**Fig. 1f&g**), generating a point-wise scalp-to-cortex distance map (**Fig. 1h**). To search efficiently for the nearest points, we use a K-dimensional tree data structure to speed up the computation process[35]. The computation took approximately 3 minutes to generate the point-wise scalp-to-cortex map, where the user could manually choose a location on the scalp surface with a mouse cursor to retrieve the value. In addition, the individual brain volume was also spatially aligned to the brain atlas (BCI-DNI) [36], with 66 regions labeled on the surface (**Fig. 1i**) using the BrainSuite registration module SVReg [37]. As a result, individual cortical surfaces were labeled accordingly (**Fig. 1j**). Finally, for each labeled region, we calculated the average value of all the point-wise scalp-to-cortex distances within the region and defined it as the SRD. Because the entire pipeline is automated as run by command lines for large datasets, certain distance thresholds were set to exclude unrealistic results caused by poor segmentation instead of manual parameter correction to each image. The whole pipeline took approximately 20 minutes for one MR image as tested on a Lenovo ThinkPad T14 with Intel i7-1355U CPU.

**Fig. 1.**
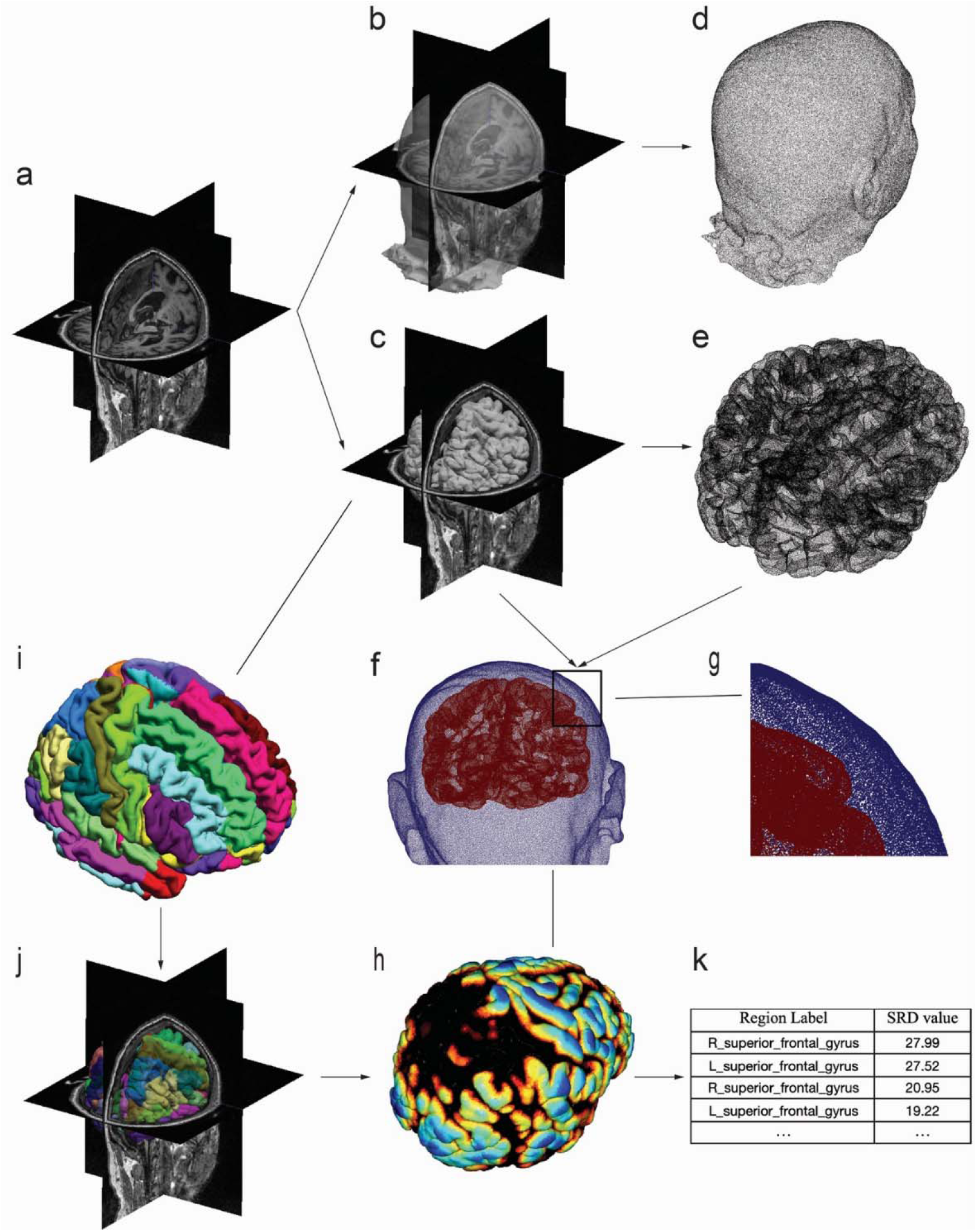
Analysis pipeline for SRD using T1W MR data. (**a**) Original T1w MRI data; (**b**) scalp; (**c**) cortex surface of the brain; (**d, e**) point clouds for the scalp surface (**d**) and pial surface (**e**); (**f, g**) zoomed-in view of the scalp and cortex points; (**h**) point-wise scalp-to-cortex distance map; (**i**) brain atlas (BCI-DNI) with labeled brain regions; (**j**) segmented regions of the cortex surface after registration with the atlas; (**k**) SRD values calculated as the average of the point-wise scalp-to-cortex distance values (in **h**) within each region (in **j**). CN, cognitively normal; AD, Alzheimer’s disease; PD, Parkinson’s disease; ASD, autism spectrum disorder. Age data are presented as the mean ± standard deviation.

### 2.3 Statistics

Statistical analyses were performed via R (version 4.5.2). Demographic and SRD variables were compared between the AD/PD/ASD group and the corresponding CN group. Age was compared via an independent sample t test. The sex distributions were compared via the chi-square test. Missing data was handled using a multiple imputation method based on Bayesian models specifically designed for large-scale neuroimaging datasets[38]. To examine region-specific SRD differences between patients and controls, analysis of covariance (ANCOVA) was performed for each brain region, with diagnostic group as the fixed factor and age and sex included as covariates. Adjusted (least-squares) means were computed to obtain covariate-adjusted group estimates. The group effect was evaluated via the t statistic associated with the group coefficient in the linear model. To account for multiple comparisons across brain regions, p values from the region-wise ANCOVAs were corrected via the false discovery rate (FDR) method. A FDR-corrected p value < 0.05 was considered statistically significant.

## 3. Results

### 3.1 Demographics

Demographic information is presented in **Table 1**. A total of 1382 participants were included in our analysis, including 313 ASD patients (15.7 ± 5.69 years, 17.6% female), 225 PD patients (61.73 +9.49 years, 64% female), 155 AD patients (74.66 ± 7.43 years, 51.6% female) and matched cognitively normal controls (CNs) in each dataset. No significant difference was found between each patient group (AD/PD/ASD) and the corresponding CN group except for sex between the ASD group and the corresponding CN group (11.5% female in the CN group from the ABIDE, p = 0.024).

### 3.4 SRD alterations in patients with ASD

The coregistered cortical regions and scalp-to-cortex distance maps of representative ASD patients and CNs are shown in **Fig. 2a&b**. The SRD alterations in ASD patients, as shown in **Fig. 2c** and **Table 2** (FDR-corrected p < 0.05), were relatively widespread. Thirty-two of the 66 tested regions (48.5%) presented significant SRD decreases in AD patients compared with those in CN, and most of them were bilateral, while no significant increase in SRD was found. Among them, more than half (18/32) of the alterations were in the frontal lobe. Notably, 12 regions were located in the orbitofrontal and adjacent gyri, including the bilateral lateral, bilateral posterior, right middle and anterior orbitofrontal gyri, the bilateral rectus and subcallosal gyri and the adjacent bilateral temporal pole, as shown by the transverse and sagittal views in **Fig. 2c**. In addition, 8 other frontal regions that spread in the dorsal direction from the bilateral superior, middle, and pars triangularis subregions of the inferior frontal gyrus to the precentral gyrus were involved. Decreased SRDs were also detected in 8 parietal regions (bilateral postcentral, superior parietal, supramarginal and angular gyri), 3 adjacent occipital regions (bilateral middle and left inferior occipital gyri) and the left middle temporal gyrus.

**Table 2.** SRD values in the ASD and corresponding CN groups.

| Region | Abbr. | Adjusted<br>mean<br>ASD | Adjusted<br>mean<br>CN | p | Corrected<br>p |
| --- | --- | --- | --- | --- | --- |
| R_superior_frontal_gyrus | SFG.R | 24.43 | 24.84 | 0.0071 | 0.0179 |
| L_superior_frontal_gyrus | SFG.L | 24.14 | 24.55 | 0.0066 | 0.0174 |
| R_middle_frontal_gyrus | MFG.R | 21.59 | 22.08 | 0.0013 | 0.0056 |
| L_middle_frontal_gyrus | MFG.L | 20.70 | 21.35 | 0.0000 | 0.0012 |
| R_pars_triangularis | TrIFG.R | 21.00 | 21.54 | 0.0006 | 0.0038 |
| L_pars_triangularis | TrIFG.L | 22.26 | 22.71 | 0.0041 | 0.0124 |
| R_pre_central_gyrus | PrG.R | 23.19 | 23.81 | 0.0007 | 0.0038 |
| L_pre_central_gyrus | PrG.L | 23.37 | 24.01 | 0.0007 | 0.0038 |
| R_gyrus_rectus | GRe.R | 21.62 | 22.85 | 0.0008 | 0.0040 |
| L_gyrus_rectus | GRe.L | 24.56 | 25.62 | 0.0006 | 0.0038 |
| R_middle_orbito_frontal_gyrus | MOrG.R | 17.97 | 18.84 | 0.0024 | 0.0087 |
| R_anterior_orbito_frontal_gyrus | AOrG.R | 17.28 | 17.96 | 0.0159 | 0.0340 |
| R_posterior_orbito_frontal_gyrus | POrG.R | 23.03 | 24.06 | 0.0025 | 0.0087 |
| L_posterior_orbito_frontal_gyrus | POrG.L | 23.42 | 24.20 | 0.0160 | 0.0340 |
| R_lateral_orbitofrontal_gyrus | LOrG.R | 17.33 | 17.91 | 0.0004 | 0.0038 |
| L_lateral_orbitofrontal_gyrus | LOrG.L | 17.14 | 17.57 | 0.0216 | 0.0446 |
| R_subcallosal_gyrus | SCG.R | 23.84 | 25.95 | 0.0105 | 0.0248 |
| L_subcallosal_gyrus | SCG.L | 24.33 | 26.55 | 0.0066 | 0.0174 |
| R_post_central_gyrus | PoG.R | 24.47 | 25.11 | 0.0016 | 0.0066 |
| L_post_central_gyrus | PoG.L | 24.30 | 24.94 | 0.0006 | 0.0038 |
| R_supramarginal_gyrus | SMG.R | 25.62 | 26.11 | 0.0049 | 0.0140 |
| L_supramarginal_gyrus | SMG.L | 25.15 | 25.80 | 0.0004 | 0.0038 |
| R_angular_gyrus | AnG.R | 22.35 | 22.84 | 0.0028 | 0.0094 |
| L_angular_gyrus | AnG.L | 20.80 | 21.63 | 0.0001 | 0.0012 |
| R_superior_parietal_gyrus | SPL.R | 22.96 | 23.82 | 0.0000 | 0.0010 |
| L_superior_parietal_gyrus | SPL.L | 23.19 | 23.90 | 0.0003 | 0.0038 |
| R_temporal_pole | tmp.R | 22.82 | 23.65 | 0.0005 | 0.0038 |
| L_temporal_pole | tmp.L | 23.12 | 23.90 | 0.0025 | 0.0087 |
| L_middle_temporal_gyrus | MTG.L | 22.12 | 22.53 | 0.0088 | 0.0216 |
| R_middle_occipital_gyrus | MOcG.R | 19.67 | 20.31 | 0.0007 | 0.0038 |
| L_middle_occipital_gyrus | MOcG.L | 17.85 | 18.45 | 0.0035 | 0.0110 |
| L_inferior_occipital_gyrus | IOG.L | 19.59 | 20.18 | 0.0125 | 0.0285 |

The listed regions show significant group differences with FDR-corrected p values < 0.05. CN, cognitively normal; ASD, autism spectrum disorder;

**Fig. 2.**
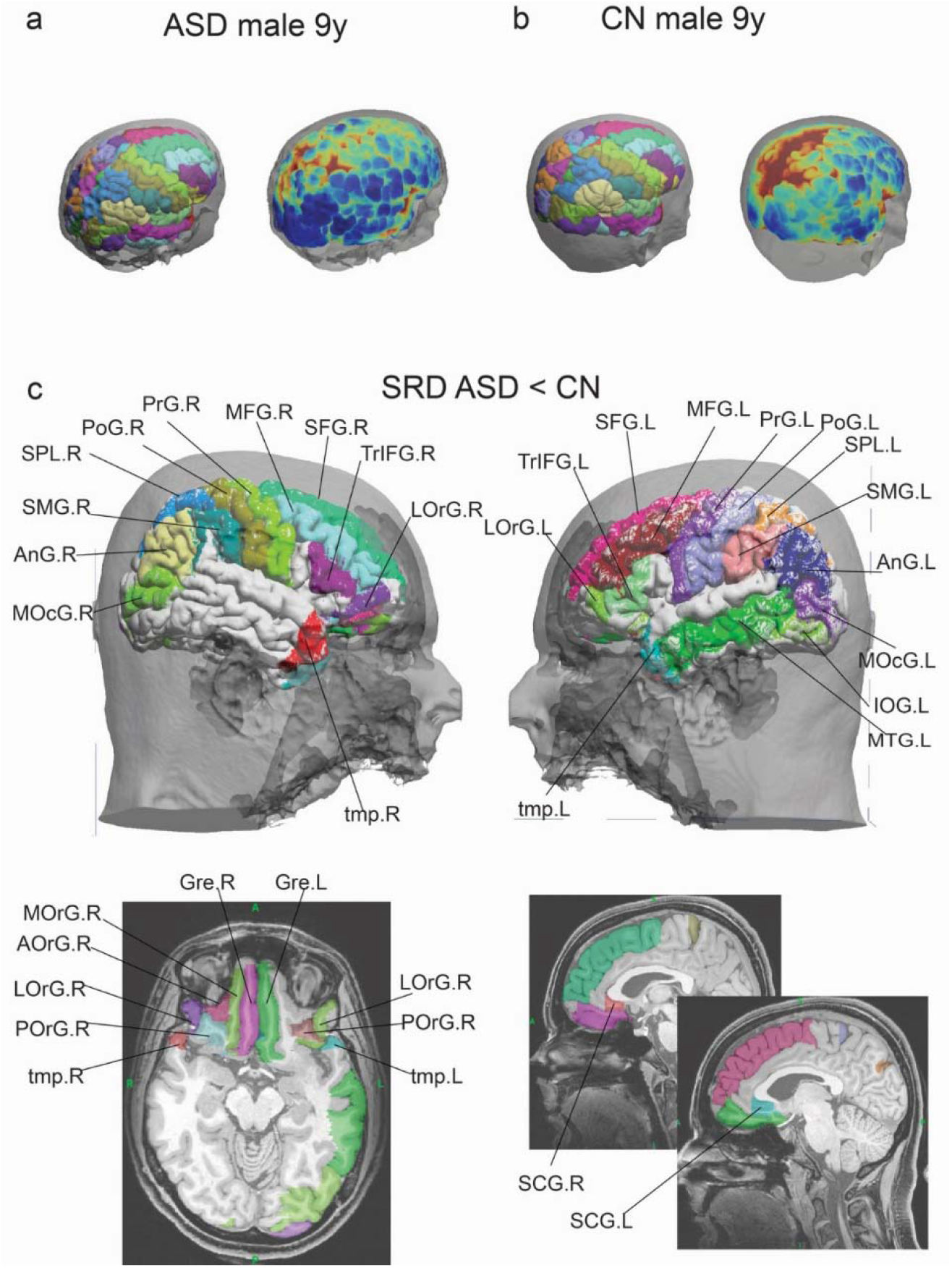
Significant SRD alterations in patients with ASD. (**a**) Coregistered brain regions and scalp-to-cortex distance map of a representative ASD; (**b**) Coregistered brain regions and scalp-to-cortex distance map of a representative CN in ABIDE; (**c**) Regions with significantly decreased SRD in the ASD group in comparison with the CN group (p<0.05 after FDR correction). The regions in **c** are shown via the BCI-DNI atlas, and different colors represent different regions. The medial regions are also illustrated in transverse and sagittal views for better illustration. SRD, scalp-to-region distance; CN, cognitively normal; ASD, autism spectrum disorder; L, left; R, right. AnG, angular gyrus; AOrG, anterior orbitofrontal gyrus; GRe, gyrus rectus; IOG, inferior occipital gyrus; LOrG, lateral orbitofrontal gyrus; MFG, middle frontal gyrus; MOcG, middle occipital gyrus; MOrG, middle orbitofrontal gyrus; MTG, middle temporal gyrus; PoG, postcentral gyrus; POrG, posterior orbitofrontal gyrus; PrG, precentral gyrus; SCG, subcallosal gyrus; SFG, superior frontal gyrus; SMG, supramarginal gyrus; SPL, superior parietal gyrus; tmp, temporal pole; TrIFG, pars triangularis subregion of the inferior frontal gyrus.

### 3.3 SRD in patients with PD

The coregistered cortical regions and scalp-to-cortex distance maps of representative PD patients and CNs are shown in **Supplementary Fig. 1a&b**. In PD patients, several regions tended to be altered, but the p values were uncorrected (uncorrected p < 0.05), as shown in **Supplementary Fig. 1c&d** and **Supplementary Table 1**. These trends included increases in SRD (**Supplementary Fig. 1c**) in the left transverse frontal and middle temporal gyri and right occipital regions involving decreases in the precuneus, cuneus and lingual gyri and SRD (**Supplementary Fig. 1d**) in the right middle temporal gyrus. However, none of these differences survived FDR correction (p<0.05).

### 3.2 SRD alterations in patients with AD

SRD alterations in AD patients are illustrated in **Fig. 3** and **Table 3** (FDR-corrected p < 0.05). The coregistered cortical regions and scalp-to-cortex distance maps of the representative AD and CN are shown in **Fig. 3a&b**. Compared with the CN, 20 of the 66 tested regions (30.3%) presented significant SRD alterations in AD patients. Significantly increased SRDs in AD patients (**Fig. 3c**) were mainly in a series of regions in the frontal lobe, including the bilateral superior, bilateral middle, inferior (bilateral pars triangularis subregion and right pars orbitalis subregion), bilateral transverse, bilateral lateral orbitofrontal, right anterior orbitofrontal frontal gyri and left precentral gyri. The bilateral superior occipital gyrus, left middle occipital gyrus and supramarginal gyrus also had greater SRD. In contrast, the bilateral parahippocampal gyrus and right fusiform gyrus, which are located in the medial temporal lobe, showed decreased SRD in AD patients (**Fig. 3d**).

**Table 3.** SRD values in the AD group and corresponding CN group.

| Region | Abbr. | Adjusted<br>mean<br>AD | Adjusted<br>mean<br>CN | p | Corrected<br>p |
| --- | --- | --- | --- | --- | --- |
| R_superior_frontal_gyrus | SFG.R | 29.11 | 28.18 | 0.0129 | 0.0426 |
| L_superior_frontal_gyrus | SFG.L | 28.73 | 27.65 | 0.0057 | 0.0308 |
| R_middle_frontal_gyrus | MFG.R | 25.31 | 23.76 | 0.0001 | 0.0024 |
| L_middle_frontal_gyrus | MFG.L | 24.84 | 23.37 | 0.0001 | 0.0024 |
| R_pars_triangularis | TrIFG.R | 25.86 | 24.70 | 0.0024 | 0.0198 |
| L_pars_triangularis | TrIFG.L | 26.09 | 25.03 | 0.0041 | 0.0300 |
| R_pars_orbitalis | OrIFG.R | 29.46 | 28.50 | 0.0048 | 0.0307 |
| L_pre_central_gyrus | PrG.L | 27.10 | 25.73 | 0.0008 | 0.0093 |
| R_transvers_frontal_gyrus | TFg.R | 23.81 | 22.52 | 0.0005 | 0.0063 |
| L_transvers_frontal_gyrus | TFg.L | 24.19 | 22.78 | 0.0001 | 0.0024 |
| R_anterior_orbito_frontal_gyrus | AOrG.R | 27.21 | 26.06 | 0.0061 | 0.0308 |
| R_lateral_orbitofrontal_gyrus | LOrG.R | 22.69 | 21.65 | 0.0109 | 0.0394 |
| L_lateral_orbitofrontal_gyrus | LOrG.L | 22.36 | 21.00 | 0.0011 | 0.0103 |
| L_supramarginal_gyrus | SMG.L | 28.99 | 27.99 | 0.0100 | 0.0394 |
| R_fusiforme_gyrus | FuG.R | 38.41 | 39.18 | 0.0110 | 0.0394 |
| R_parahippocampal_gyrus | PHG.R | 49.02 | 50.57 | 0.0051 | 0.0307 |
| L_parahippocampal_gyrus | PHG.L | 50.04 | 52.38 | 0.0000 | 0.0018 |
| R_superior_occipital_gyrus | SOG.R | 20.13 | 19.06 | 0.0070 | 0.0329 |
| L_superior_occipital_gyrus | SOG.L | 18.55 | 17.51 | 0.0113 | 0.0394 |
| L_middle_occipital_gyrus | MOcG.L | 21.15 | 20.08 | 0.0098 | 0.0394 |

The listed regions show significant group differences with FDR-corrected p values < 0.05. CN, cognitively normal; AD, Alzheimer's disease;

**Fig. 3.**
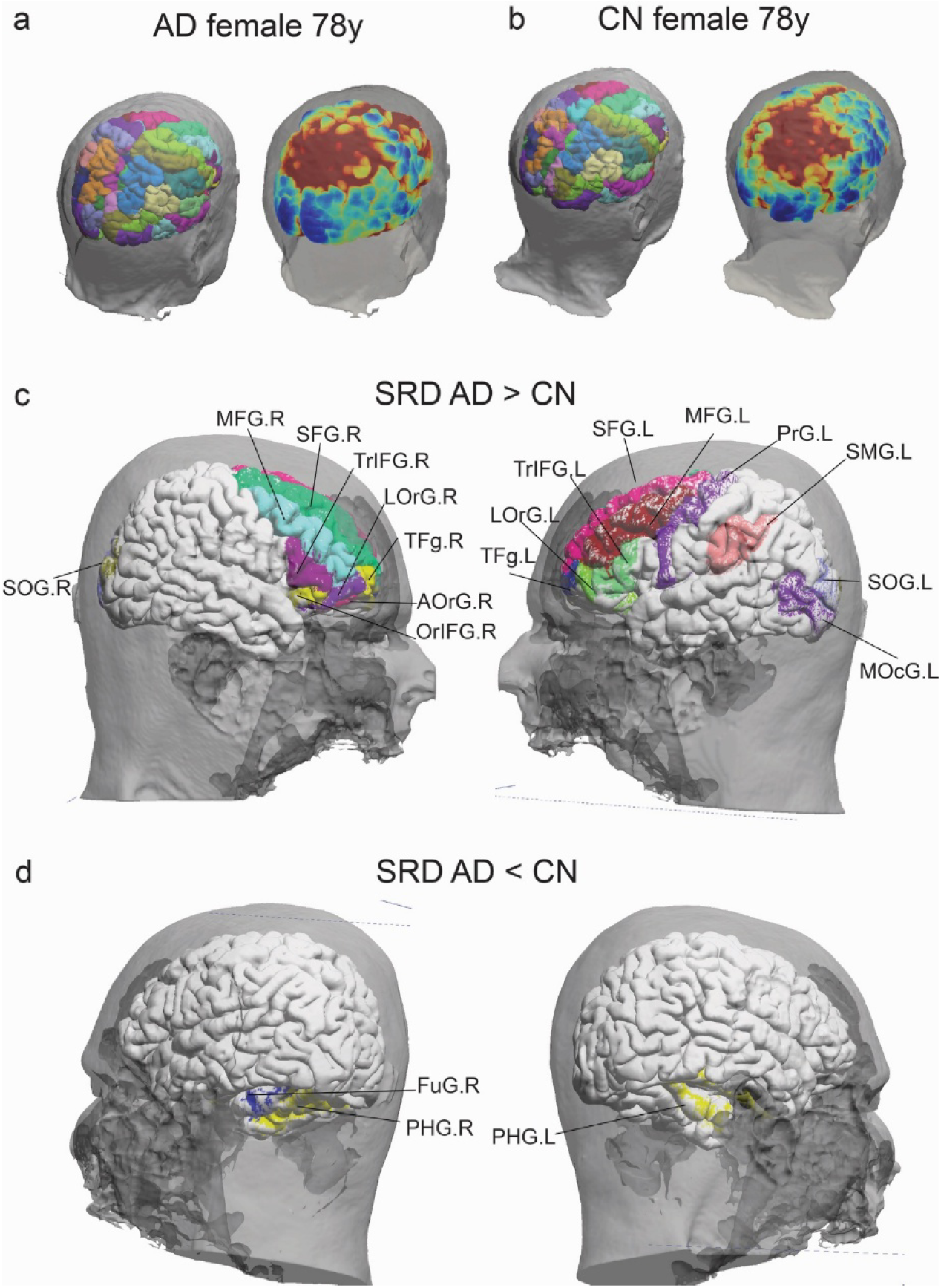
Significant SRD alterations in patients with AD. (**a**) Coregistered brain regions and scalp-to-cortex distance map of a representative AD; (**b**) Coregistered brain regions and scalp-to-cortex distance map of and a representative CN in the ADNI; (**c**) Regions with significantly increased SRD in the AD group in comparison with the CN group; (**d**) Regions with significantly decreased SRD in the AD group in comparison with the CN group. Regions with significant alterations (p<0.05 after FDR correction) in **c&d** are shown via the BCI-DNI atlas, and different colors represent different regions. SRD, scalp-to-region distance; CN, cognitively normal; AD, Alzheimer’s disease; L, left; R, right. AOrG, anterior orbitofrontal gyrus; FuG, fusiform gyrus; LOrG, lateral orbitofrontal gyrus; MFG, middle frontal gyrus; MOcG, middle occipital gyrus; OrlFG, pars orbitalis subregion of the inferior frontal gyrus; PHG, parahippocampal gyrus; PrG, precentral gyrus; SFG, superior frontal gyrus; SMG, supramarginal gyrus; SOG, superior occipital gyrus; TFg, transverse frontal gyrus; TrIFG, pars triangularis subregion of the inferior frontal gyrus.

A schematic illustration of the disease-specific significant SRD alterations in the ASD and AD groups is shown in **Fig. 4**. The two diseases clearly exhibit different SRD patterns. The SRD decreases in the ASD patients ranged from -0.41mm to -2.22mm, whereas those in the AD patients ranged mainly showed SRD increases from 0.94mm to 1.55mm and several SRD decreases ranged from -0.76mm to -2.33mm.

**Fig. 4.**
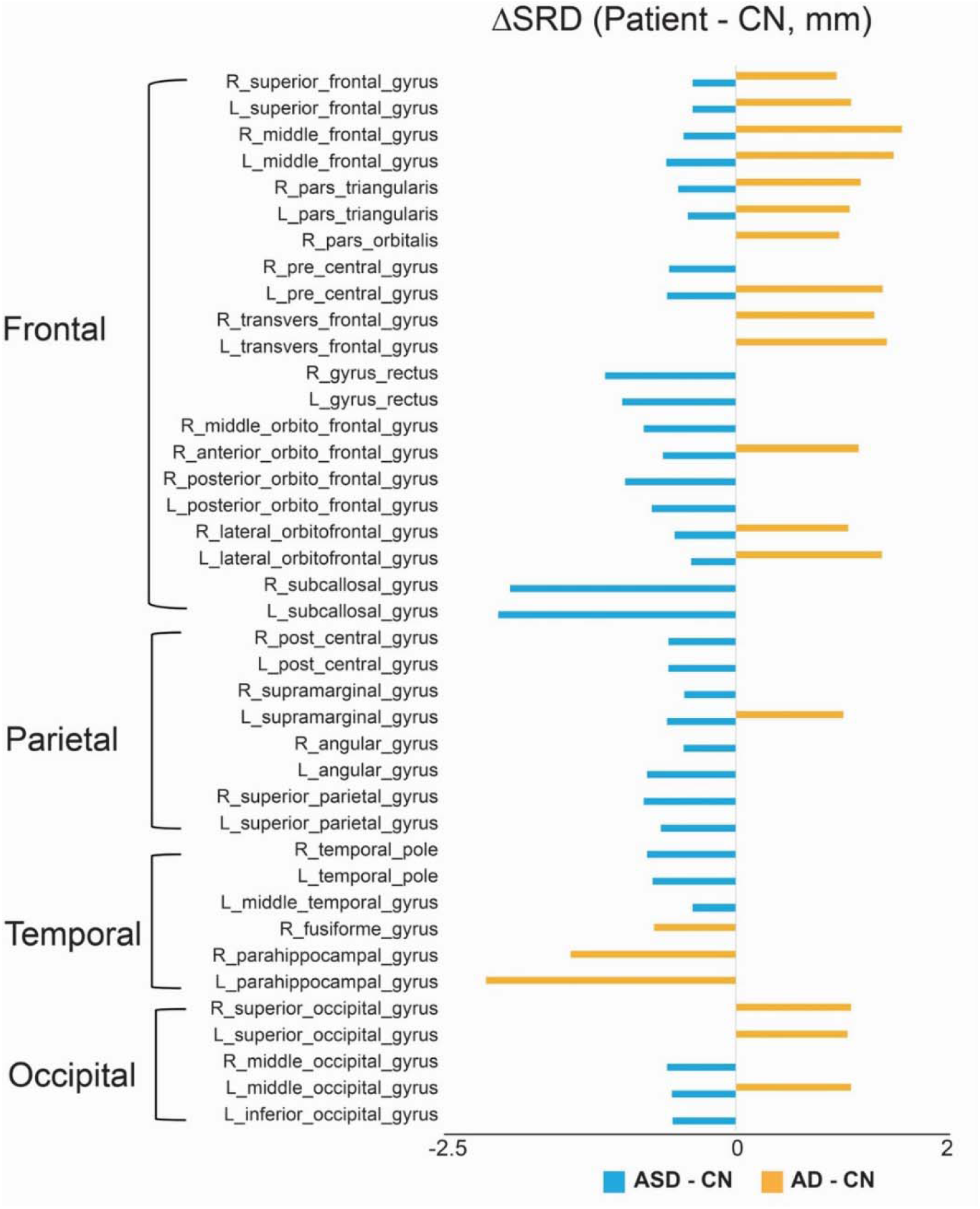
Bar plots showing disease-specific SRD alterations in ASD and AD groups. The length of the bars represents the difference in SRD (mm) between patients and healthy controls. SRD, scalp-to-region distance; CN, cognitively normal; ASD, autism spectrum disorder; AD, Alzheimer’s disease.

## 4 Discussion

In this study, we developed an automatic open-source pipeline that enabled efficient regional SRD analysis. Uins this pipeline, we performed automatic analysis of SRD in a large sample (N = 1382) and reported different SRD alteration patterns in three different neurological disorders, including one neurodevelopmental disease (ASD) and two neurodegenerative diseases (PD and AD). Among the three diseases, ASD patients showed widespread SRD decreases across the brain with prominent involvement of the frontal lobe, especially the orbitofrontal cortex and adjacent regions. Conversely, AD patients presented significantly increased SRD values in various regions of the frontal gyrus. No significant SRD alteration was found in PD patients after correction.

Elucidating alterations in scalp-to-region distance in patients with brain diseases could provide an essential reference for the development of novel neurostimulation targets and treatment planning. However, such investigation is compromised in previous studies [17, 20-24] by methodological drawbacks including manual selection of interested brain or scalp pixel/voxel, subjective delineation of measuring lines\boxes and limited number of brain areas and sample size consequently. The scalp-to-cortex distance calculation pipeline proposed in our previous study was able to derive individual maps with numerous points spread across the entire cortex surface[26]. In this study, we further developed a pipeline with a new function module, the automatic calculation of SRD, which is not available in any current toolboxes, and applied it to various brain disorders with a large sample size.

Our results revealed different SRD alteration patterns in patients with ASD, PD and AD. Intriguingly, the ASD and AD patients showed substantially divergent, and even opposite SRD patterns in some regions. The mechanisms underlying SRD alterations in patients may be multifaceted. Abnormalities in structures along the trajectory from the scalp-to-cortex of the brain may contribute to the variability of SRD, including cortical geometry, gray matter volume, skull thickness, etc.

ASD is a heterogeneous neurodevelopmental disorder characterized by deficiencies in social perception and abilities and is associated with a variety of environmental and genetic factors [39]. To date, pharmaceutical interventions for alleviating the fundamental symptoms of ASD have been limited, and behavioral therapies and education have been shown to have only minor-to-moderate effects. Neurostimulation treatments, including TMS[6] and tDCS[2], show some potential but exhibit marked heterogeneity, posing a challenge to the selection of treatment targets and parameters. Interestingly, in addition to earlier studies revealing increased cortical gyrification[40] and overall greater subcortical gray matter volume [41] in ASD patients, recent large-sample ASD studies have also reported greater cortical thickness involving the bilateral inferior frontal and prefrontal cortex (491 individuals with ASD) [42] and increased cortical thickness in the frontal cortex, with a peak around adolescence across the lifespan (1,571 patients with ASD)[43]. Specifically, the increase in the orbitofrontal cortex and anterior cingulate cortex in high-compared with medium-aggression ASD patients[44] indicates a functional association among these brain regions. The expansion of these different brain structures, especially the frontal cortex, may collectively contribute to the decreased SRD in ASD as revealed by our results.

PD is a progressive neurodegenerative illness with both motor and nonmotor symptoms [45]. SRD values were not significantly altered in PD patients, according to our results. A previous study based on 47 early-stage PD patients reported increased scalp-to-cortex distance in the left DLPFC, the most widely used target in TMS treatment[23]. Such discrepancies may arise from differences in point/area selection, distance calculation methods and population characteristics. Notably, the well-validated stimulation targets associated with the brain mechanism of PD motor symptoms are mainly the deep nuclei, including the subthalamic nucleus (STN) and globus pallidus internus (GPi)[1], both of which have much smaller volumes than the whole head space does. In addition, the nonmotor symptoms of PD are heterogeneous and involve a series of cognitive and neuropsychiatric symptoms that respond differently to various stimulation treatments[46], weakening the significance of alterations at the group level.

AD and its prodromal stage mild cognitive impairment (MCI) constitute a disease continuum characterized by progressive cognitive decline and memory loss [47]. Neuropathological studies shows that AD-associated neurofibrillary pathology begins in the transentorhinal region, entorhinal region and hippocampus; spreads to the fusiform and temporal neurocortex; and finally extends to the frontal, parietal and occipital neocortex[48]. Positron emission tomography using tau tracers have provided visualizing of similar pattern in AD patients [49-51]. Interestingly, our results of AD patients revealed decreased SRD in the early AD staging area, involving the bilateral parahippocampal gyrus (with the entorhinal cortex as its anterior portion and adjacent to the deep hippocampus structure) and right fusiform gyrus, which may be helpful for planning the neurostimulation of AD patients and further studying the complex mechanism involved. Conversely, a variety of frontal regions in AD patients showed increased SRD, which is also markedly different from that observed in patients with ASD. A recent study analyzed and reported a particular increase in the distance from the scalp-to-cortex in two frontal regions, the DLPFC and the primary motor cortex, in 43 dementia patients[24]. Our results not only confirmed SRD alterations in the two brain regions but also comprehensively illustrated SRD alterations across whole-brain surface regions in AD patients.

There are several limitations in this study. 1) The reasons behind the SRD alterations are discussed but not analyzed in this study because complex factors along the trajectory from the scalp to the cortex of the brain may contribute to the variability of SRD, whose relationships need to be further studied in future studies. 2) Although this study utilized a large dataset (N=1382), the findings need to be validated using more multi-center data before clinical practice.

## Conclusion

In this study, we developed an automatic SRD calculation pipeline and applied it to ASD, PD and AD, revealing different SRD alteration patterns. These tools and findings are potentially helpful for planning the neurostimulation of patients with brain diseases in clinical practice.

## Author Contributions

RN designed the study; JZ wrote the code; JZ, LY, JW, and HH performed the analysis; LY and RN interpreted the data; LY, JZ and RN wrote the draft. All the authors read and approved the final manuscript.

## Acknowledgments

ADNI: Data collection and sharing for this project was funded by the ADNI (National Institutes of Health Grant U01 AG024904) and DOD ADNI (Department of Defense award number W81XWH-12-2-0012). The ADNI is funded by the National Institute on Aging, the National Institute of Biomedical Imaging and Bioengineering, and through generous contributions from the following: AbbVie, Alzheimer’s Association; Alzheimer’s Drug Discovery Foundation; Araclon Biotech; BioClinica, Inc.; Biogen; Bristol-Myers Squibb Company; CereSpir, Inc.; Cogstate; Eisai Inc.; Elan Pharmaceuticals, Inc.; Eli Lilly and Company; EuroImmun; F. Hoffmann-La Roche Ltd. and its affiliated company Genentech, Inc.; Fujirebio; GE Healthcare; IXICO Ltd.; Janssen Alzheimer Immunotherapy Research & Development, LLC.; Johnson & Johnson Pharmaceutical Research & Development LLC.; Lumosity; Lundbeck; Merck & Co., Inc.; Meso Scale Diagnostics, LLC.; NeuroRx Research; Neurotrack Technologies; Novartis Pharmaceuticals Corporation; Pfizer Inc.; Piramal Imaging; Servier; Takeda Pharmaceutical Company; and Transition Therapeutics. The Canadian Institutes of Health Research is providing funds to support ADNI clinical sites in Canada. Private sector contributions are facilitated by the Foundation for the National Institutes of Health (www.fnih.org). The grantee organization is the Northern California Institute for Research and Education, and the study is coordinated by the Alzheimer’s Therapeutic Research Institute at the University of Southern California. ADNI data are disseminated by the Laboratory for Neuro Imaging at the University of Southern California.

PPMI: a public□private partnership, is funded by the Michael J. Fox Foundation for Parkinson’s Research and funding partners, including 4D Pharma, AbbVie, AcureX, Allergan, Amathus Therapeutics, Aligning Science Across Parkinson’s, AskBio, Avid Radiopharmaceuticals, BIAL, BioArctic, Biogen, Biohaven, BioLegend, BlueRock Therapeutics, Bristol-Myers Squibb, Calico Labs, Capsida Biotherapeutics, Celgene, Cerevel Therapeutics, Coave Therapeutics, DaCapo Brainscience, Denali, Edmond J. Safra Foundation, Eli Lilly, Gain Therapeutics, GE HealthCare, Genentech, GSK, Golub Capital, Handl Therapeutics, Insitro, Jazz Pharmaceuticals, Johnson & Johnson Innovative Medicine, Lundbeck, Merck, Meso Scale Discovery, Mission Therapeutics, Neurocrine Biosciences, Neuron23, Neuropore, Pfizer, Piramal, Prevail Therapeutics, Roche, Sanofi, Servier, Sun Pharma Advanced Research Company, Takeda, Teva, UCB, Vanqua Bio, Verily, Voyager Therapeutics, the Weston Family Foundation and Yumanity Therapeutics.

ABIDE: ASD data were provided by the Autism Brain Imaging Data Exchange (ABIDE) and contributors. The ABIDE data used in the preparation of this manuscript were supported by the ABIDE funding resources listed at http://fcon_1000.projects.nitrc.org/indi/abide/. ABIDE’s primary support for the work by Adriana Di Martino was provided by the NIMH (K23 MH087770) and the Leon Levy Foundation. Primary support for the work by Michael P. Milham and the INDI team was provided by gifts from Joseph P. Healy and the Stavros Niarchos Foundation to the Child Mind Institute.

