## Supplemental files for "Divergent scalp-to-region distance alteration patterns in autism spectrum disorders, Parkinson’s disease and Alzheimer’s disease"

**Supplementary materials**

**Supplementary Table 1 SRD values in the PD and corresponding CN groups**

| Region |  | Abbr. | Adjusted mean  PD | Adjusted mean  CN | p | Corrected  p |
| --- | --- | --- | --- | --- | --- | --- |
| L_transvers_frontal_gyrus | | TFg.L | 22.74 | 21.64 | 0.0259 | 0.4698 |
| R_pre_cuneus | | Pcu.R | 39.91 | 39.17 | 0.0383 | 0.4698 |
| R_middle_temporal_gyrus | | MTG.R | 22.81 | 23.21 | 0.0280 | 0.4698 |
| L_middle_temporal_gyrus | | MTG.L | 23.62 | 23.24 | 0.0427 | 0.4698 |
| R_lingual_gyrus | | LiG.R | 41.26 | 40.43 | 0.0120 | 0.4698 |
| R_cuneus |  | Cun.R | 32.41 | 31.75 | 0.0417 | 0.4698 |

The listed regions show a trend toward group differences with uncorrected p values < 0.05, which are not significant after FDR correction. SRD, scalp-to-region distance; CN, cognitively normal; PD, Parkinson’s disease;

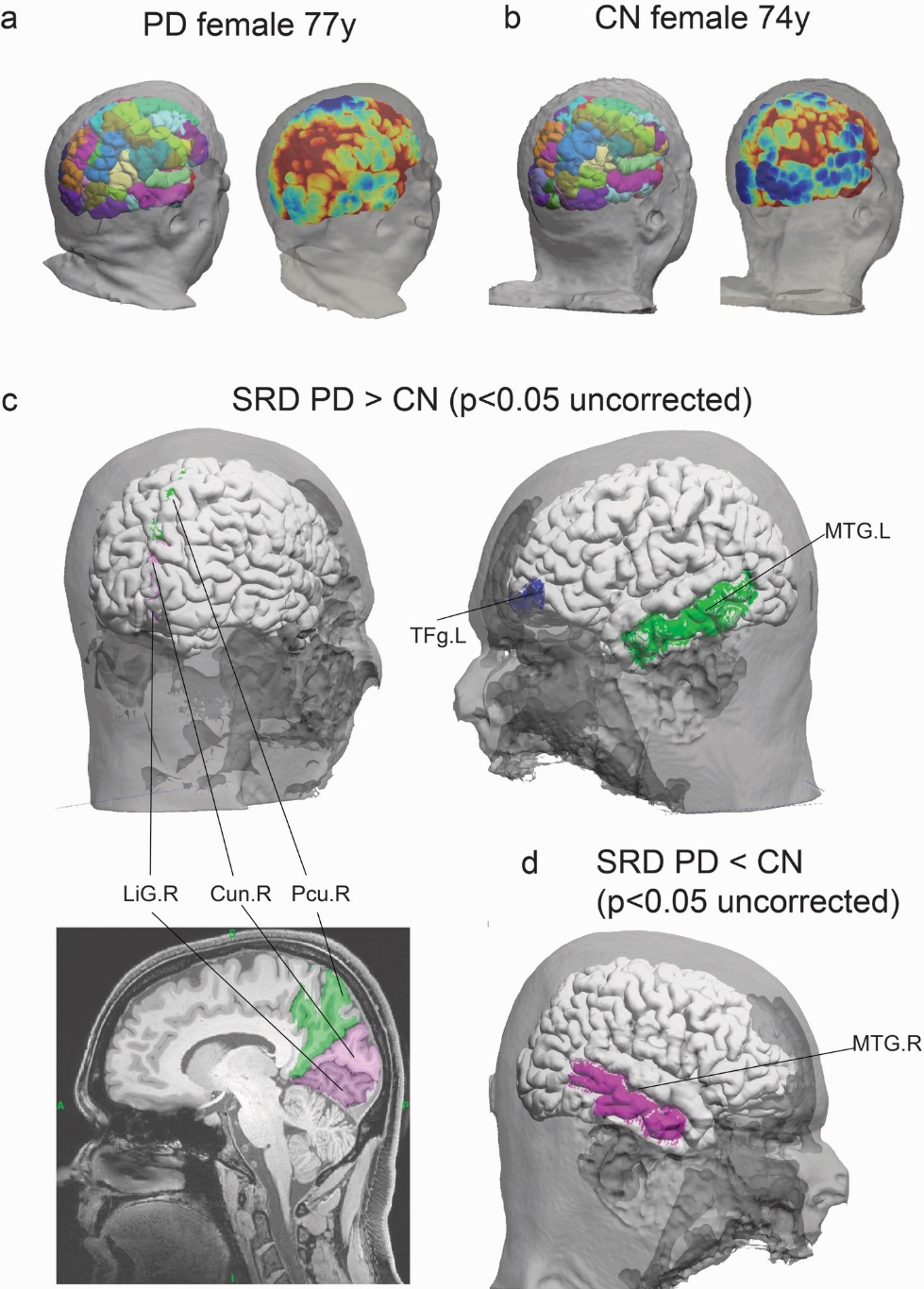

**Supplementary Fig. 1 SRD trends in patients with PD.** (**a**) Coregistered brain regions and scalp-to-cortex distance map of a representative PD; (**b**) Coregistered brain regions and scalp-to-cortex distance map of a representative CN in the PPMI; (**c**) Regions with a trend toward increasing SRD in the PD group in comparison with the CN group; (**d**) Regions with a trend toward decreasing SRD in the PD group in comparison with the CN group. Notably, these trends of differences were found with uncorrected p<0.05, and none of them survived with FDR-corrected p<0.05. The regions in **c**&**d** are shown via the BCI-DNI atlas, and different colors represent different regions. The medial regions are also illustrated in the sagittal view for better illustration. SRD, scalp-to-region distance; CN, cognitively normal; AD, Alzheimer’s disease; L, left; R, right. Cun, Cuneus; LiG, lingual gyrus; MTG, middle temporal gyrus; Pcu, precuneus; TFg, transverse frontal gyrus.
